# Gene expression changes in long-term memory unlikely to replicate in the long term

**DOI:** 10.1101/2024.07.22.604349

**Authors:** Eran A. Mukamel, Zhaoxia Yu

## Abstract

Identifying the cellular effects of memory-forming experiences on neurons which enable subsequent memory recall is a fundamental aim of neuroscience. The search for the engram could benefit from single cell RNA sequencing, which can estimate the mRNA expression of all genes in large samples of individual brain cells from animals exposed to specific experiences. A recent study used spatial transcriptomics and single cell RNA-sequencing to identify “transcriptional signatures in subpopulations of neurons and astrocytes that were memory-specific and persisted for weeks”^1^. However, because the authors did not account for multiple statistical comparisons^2^ and instead used an “unadjusted” threshold for statistical significance, the reported findings are likely dominated by false positives^3^. Moreover, the statistical analysis treated individual cells as independent samples without accounting for correlations across cells derived from the same biological tissue sample. Reanalysis of the study’s data using appropriate, widely accepted statistical procedures, identifies no significant differentially expressed genes. This suggests the data do not support the author’s claim to have identified cell type-specific transcriptional signatures of memory in the mouse basolateral amygdala.

## MAIN TEXT

The risk of false positives due to multiple comparisons in large-scale genomic studies has been recognized for many years^2^. When testing the effect of a treatment on thousands of genes, around 5% of the tested genes are expected to pass an unadjusted significance threshold (p<0.05) even in the absence of any true effect. In the current study, the authors tested 3,350 genes for a significant difference of expression due to fear and recall experiences within a specific group of activated “engram” neurons (TRAPed BlaIn.Gpr88 neurons; Fig. 2e of Sun et al.). They applied a nominal (unadjusted) p-value threshold (p<0.05) as well as post-hoc criteria based on effect size (fold-change > 1.75) and specificity for the fear/recall experience. They reported 32 genes associated with remote memory in the *Gpr88* expressing inhibitory neurons, and 107 total genes passing “stringent criteria” across 6 types of neurons. Notably, the unadjusted threshold they applied would be expected to produce ∼160 false positives (5% of 3,350) in a single cell type, raising the question of whether the reported effects of memory are statistically valid.

We focus on the main finding reported in the paper, which is that a specific population of GABAergic “engram” neurons (TRAPed, defined by expression of tdT) exhibited a pattern of up- and down-regulation of several hundred genes that was specifically induced by fear and recall (FR) experiences (see Methods). The authors identified these genes by comparing the expression of genes in FR mice with control mice that did not experience fear conditioning (no-fear, NF). Here we focused on the main analysis of a population of inhibitory neurons in the BLA expressing Gpr88 (BlaIn.Gpr88), but our findings apply equally to the other cell types which were analyzed in the same way.

We first verified that, consistent with the original paper, analyzing the data with no adjustment for multiple comparisons yields 253 nominally significant genes (p<0.05), of which 230 also had absolute fold-change >1.75 (Fig. 1a). The 107 strictly filtered genes described by the authors are presumably a subset of these genes which result from applying further criteria, such as excluding genes which were also differentially expressed in non-TRAPed (tdT–) neurons. This confirms that we correctly interpreted the dataset and successfully replicated the reported analysis.

**Figure 1:**
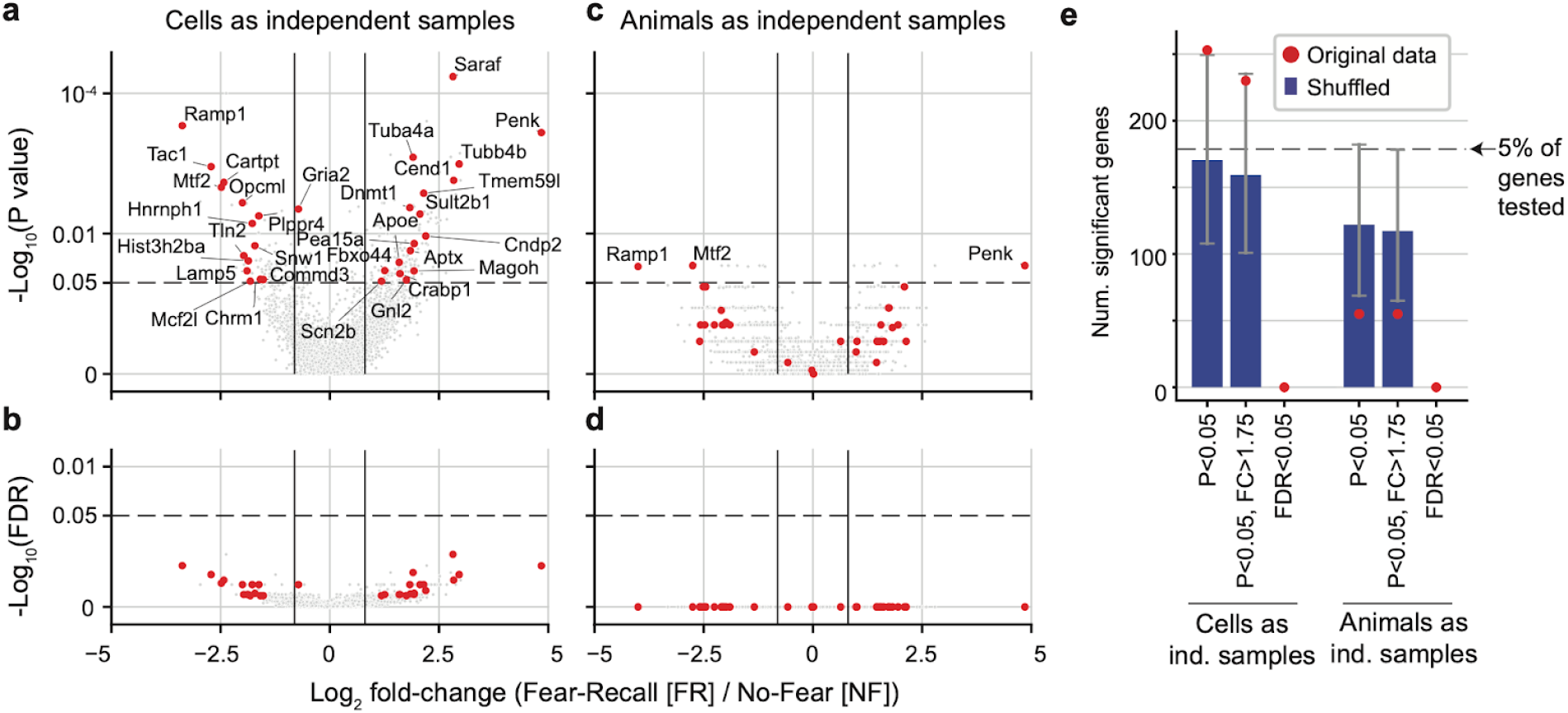
Failing to adjust for multiple comparisons leads to spurious detection of differential expression. **a-d**, Volcano plots showing the difference in mRNA expression in TRAPed BLA.Int.Gpr88 cells between mice exposed to fear and recall (FR) vs. no-fear control (NF), as in Fig. 2e of the orignal paper^1^. The x-axis shows the fold-change, and the y-axis shows the statistical significance assessed by unadjusted p-value (a,c) or using the false discovery rate (FDR) adjusted p-value (b,d) (two-sided Mann-Whitney U rank-based test). Grey dots show all tested genes, and red dots show the significant genes identified in Fig. 2e of the original paper. **a**,**b** Analysis treating individual cells as independent samples (n=70 cells). **c**,**d** Analysis using pseudobulk expression profiles from each animal as an independent sample (n=11 animals; 4 FR, 7 NF). **e**, Number of significant genes using the original data (red) or after shuffling the treatment labels (FR, NF) across cells (blue bars, 1000 shuffles; error bar shows 25th-75th percentile range).

The standard practice for controlling false positives in large-scale genomic studies is the false discovery rate (FDR), which aims to control the expected proportion of false positives among the detected significant genes. In practice, FDR-adjusted p-values are often computed to account for multiple comparisons^2,4^. Indeed, a closely related paper from the same laboratories used FDR correction to identify memory-related gene expression changes in the medial prefrontal cortext^5^. Using adjusted p-values at a specified FDR of 0.05 ensures that no more than ∼5% of the reported significant results will be false discoveries. When we applied the Benjamini-Hochberg FDR procedure, we found no genes with FDR<0.05 or <0.10; the smallest FDR was 0.18 (Fig. 1b).

A second major problem with the statistical analysis is the treatment of individual cells as independent samples, without accounting for the correlation between cells derived from the same animal. Differences among individuals are a critical source of variability in biological data^6,7^. By treating single cells as independent samples, the statistical analysis ignores the within-sample dependence and leads to overconfident results. As a consequence, the findings will have poor generalizability to other mice treated in the same way, failing to demonstrate *bona fide* effects of remote memory on engram neurons. The most reliable approach for analyzing differential expression with single cell RNA-seq data handles inter-individual variability by using pseudobulk samples that combine data from all cells observed from each biological sample^8^. Alternatively, group differences in single cell RNA-seq data can be analyzed using mixed models that include random effects to account for individual variability^7,9^.

We reanalyzed the data using pseudobulk expression profiles (n=11 mice) rather than individual cells (n=70) as the unit of analysis (Fig. 1c,d). This analysis identified only 21 genes with a nominally significant (unadjusted) p value (p<0.05). The FDR adjusted p-values for all the genes were 1.

The likelihood that most of the observed expression changes are spurious false positives can be directly demonstrated by analyzing shuffled data. We randomly permuted the treatment labels (NF, FR) of the 70 cells 1,000 times, and applied each method of differential expression analysis to each permutation of the data. The number of genes passing the author’s threshold (p<0.05) for the shuffled data (170±75 mean ± s.d.) was close to the expected number (5% of 3,574 genes = 178) (Fig. 1e). Notably, 12.3% of shuffles resulted in a larger number of nominally significant genes than the original data (253). By contrast, the FDR adjusted tests led to no false positives across the shuffles, as expected.

Progress in understanding the relationship between cognitive memory and experience-dependent changes in brain gene expression requires rigor and care in the analysis of complex single-cell transcriptomic data. By neglecting best practices such as adjustment for multiple comparisons and accounting for dependence due to multiple cells from the same animal, research based on RNA sequencing risks an accumulation of irreproducible findings^3,7^.

## Methods

Processed gene expression data from the original study were downloaded from NCBI/GEO at accession GSE246147. The identity of the 70 amygdala cells which were annotated as “TRAPed BLA.Int.Gpr88” (i.e. expressing tdT and Gpr88) was kindly provided by Wenfei Sun. The identity of the animal from which each cell derived was extracted from the first three characters of the cell id. The cells were unevenly distributed across animals, with 39/70 cells coming from two FR animals, while 7 of the 11 animals contributed ≤3 cells each.

We tested differences in expression using a Mann-Whitney U rank test, implemented in the Python packaged scipy.stats.mannwhitneyu. We included 3,574 genes which had normalized expression (counts per million, CPM) >600, in order to approximately match the analysis in Fig. 2e of the original study. Note that Fig. 2e of Sun et al. states “Total = 3,350 DEGs,” but we interpret this as referring to the number of genes tested rather than the number of significant differentially expressed genes. We calculated false discovery rate (FDR)-adjusted p-values using the Benjamini-Hochberg method^4^ implemented in Python statsmodels.stats.multitest.fdrcorrection. To assess the rate of false positives, we randomly permuted the treatment label (NF and FR) of the 70 cells and repeated the Mann-Whitney U-test and FDR correction. We repeated the random shuffling 1,000 times to assess the mean and variability of the number of nominally significant genes under the null hypothesis.

A notebook reproducing the analyses presented here is available at https://github.com/mukamel-lab/SunQuake_Nature2024_Reanalysis and will be deposited at Zenodo upon publication.

